# Stepwise increase in plasmodesmata during C_4_ evolution in *Flaveria*

**DOI:** 10.1101/2025.04.18.649591

**Authors:** Kornelija Aleksejeva, Tina B. Schreier

## Abstract

Plant cells are interconnected via plasmodesmata that enable the exchange of mobile molecules such as metabolites between neighbouring cells. In leaves of plants that conduct C_4_ photosynthesis, carbon fixation is separated between mesophyll (M) and bundle sheath (BS) cells, and the shuttling of metabolites between these cells is thought to be crucial for running the carbon concentrating mechanism. Higher numbers of plasmodesmata for metabolite exchange between M and BS cells have been observed in multiple lineages of C_4_ plants, but how this increased cell-to-cell connectivity developed in the context of C_4_ evolution is not understood. Here, we examined plasmodesmata in *Flaveria* species, including C_3_, C_3_-C_4_ intermediate, C_4_-like and C_4_ species to represent an evolutionary gradient. Using electron microscopy, we found two distinct stepwise increases in plasmodesmata frequency along this gradient. C_3_-C_4_ intermediate species had ∼3 fold more plasmodesmata than C_3_ species, and C_4_-like and C_4_ species had >6-fold increases relative to the C_3_ species. In the first step between C_3_ and C_3_-C_4_ intermediate species, plasmodesmata numbers were higher in the intermediate species, but equal in frequency between M-M and M-BS cell interfaces. However, in a second step between C_3_-C_4_ intermediate and C_4_-like/C_4_ species, the increase in plasmodesmata formation was predominantly observed at the M-BS cell interface, where C_4_/C_3_ acid (malate/alanine) shuttles are active. The higher physical cell-to-cell connectivity along the evolutionary gradient occurred without significant changes in BS cell size, but coincided with the formation of increased chloroplast area/coverage in BS cells of C_4_ species. Thus, we propose that high cell-to-cell connectivity may facilitate “C_2_ metabolism” in the C_3_-C_4_ intermediate species, which also requires metabolite transfer between cell types (e.g. glycine shuttle), but full polar enrichment of plasmodesmata at the M-BS interface accompanies C_4_ metabolism. Overall, our work provides new insight into the stepwise development of cell-to-cell connectivity along the evolutionary trajectory of C_4_ photosynthesis.

## Introduction

Cell-to-cell connectivity is vital for multicellular life, particularly in plants, where cells are immobilised within rigid cell walls. Neighbouring plant cells are connected via plasmodesmata that traverse the shared cell wall. Plasmodesmata are highly complex structures that are typically 50-100 nm in diameter, and consist of a desmotubule (an endoplasmic reticulum (ER)-derived structure that connects the ER of adjacent cells), and a cytoplasmic sleeve that enables symplastic intercellular connectivity (Tee & Faulkner, 2024; Bayer & Benitez-Alfonso, 2024). The cytoplasmic sleeve facilitates the exchange of metabolites and molecules, including hormones, peptides, and proteins between adjacent cells (Roberts & Oparka, 2003; Burch-Smith & Zambryski, 2012; Li *et al*., 2021). Plasmodesmata are therefore crucial for the intercellular communication required for coordinating plant metabolism and development, as well as responses to abiotic and biotic stresses (Otero *et al*., 2016; Tee & Faulkner, 2024; Bayer & Benitez-Alfonso, 2024).

Plasmodesmata are especially important for metabolic pathways that occur across different cell types, such as C_4_ photosynthesis which typically runs across connected mesophyll (M) and bundle sheath (BS) cells. C_4_ photosynthesis is a more efficient type of photosynthesis that evolved independently around 24–35 million years ago in over 62 different plant lineages from the ancestral C_3_ pathway (Sage, 2004; Sage *et al*., 2011). In C_3_ plants, CO_2_ is captured by <u>R</u>ib<u>u</u>lose-1,5-<u>Bis</u>phosphate <u>C</u>arboxylase/<u>O</u>xygenase (RuBisCO) in mesophyll (M) cells. However, due to its oxygenase activity, RuBisCO can also fix O_2_ instead of CO_2_, leading to the production of toxic 2-phosphoglycolate (2-PG). The regeneration of 2-PG back to the carbon acceptor ribuolase-1,5 bisphosphate (RuBP) is known as photorespiration, and is an energetically costly process (Walker *et al*., 2016). C_4_ plants have evolved a carbon-concentrating mechanism that limits photorespiration, allowing more efficient carbon fixation in hot and dry climates. In C_4_ plants, CO_2_ is initially fixed in the M cells by <u>p</u>hospho*<u>e</u>nol*<u>p</u>yruvate <u>c</u>arboxylase (PEPC) into a C_4_ acid, which is subsequently transferred to the neighbouring BS cell, where it is decarboxylated to a C_3_ acid to release CO_2_. This shuttle builds high relative CO_2_ concentrations in the BS cell for refixation by RuBisCO. The C_3_ acid (pyruvate or alanine) is then transferred/shuttled back to the M cells to restart the C_4_ cycle. Thus, the efficient operation of C_4_ photosynthesis depends on the efficient metabolic flux between M and BS cells to enable this cyclic exchange of molecules (Osmond, 1971; Hatch, 1987).

C_4_ plants have specialised Kranz leaf anatomy that facilitates C_4_ metabolism. Kranz anatomy increases contact between M and BS cells through higher vein density, a concentric arrangement of BS and M cells around veins, enlarged BS cells, and enhanced intercellular connectivity between M and BS cells via numerous plasmodesmata (Sedelnikova *et al*., 2018). Early reports observed a higher number of plasmodesmata at the M-BS cell interface of C_4_ grasses compared to C_3_ grasses, and to other cell interfaces within C_4_ species (Evert *et al*., 1977; Botha, 1992; Botha *et al*., 1993). More recent studies have demonstrated a 6-to-13-fold higher plasmodesmal frequency specifically at the M-BS interface in C_4_ species relative to C_3_ relatives, in both dicotyledonous and monocotyledonous lineages (Danila *et al*., 2016, 2018, 2019; Schreier *et al*., 2024). This enhanced plasmodesmal frequency is thought to enable greater flux of C_4_ metabolites between the two cell types via passive diffusion (Osmond, 1971; Stitt & Heldt, 1985; Danila *et al*., 2016; Schreier *et al*., 2024).

Although plasmodesmata are essential for metabolite exchange in C_4_ photosynthesis, their role in the evolutionary emergence of C_4_ traits has not been systematically examined. The evolution of C_4_ photosynthesis is thought to occur by the stepwise acquisition of C_4_ Kranz anatomy traits and there are many C_3_-C_4_ intermediate species reported in both monocots and dicots (Monson & Moore, 1989; Mercado & Studer, 2022). These C_3_-C_4_ intermediate species appear to represent an evolutionary stable state in some groups (Walsh *et al*., 2023), and are found in lineages where no C_4_ species have yet been identified, for instance *Moricandia* species within the Brassicaceae family (Schlüter *et al*., 2017, 2023). Genera that contain both intermediate C_3_-C_4_ and C_4_ species are a useful resource for studying the genetic, biochemical and anatomical adaptations involved in the evolution of C_4_ photosynthesis (*Flaveria* species: (Ku *et al*., 1983; Gowik *et al*., 2011; Schulze *et al*., 2013; Kümpers *et al*., 2017); *Steinchisma* species: (Khoshravesh *et al*., 2016); *Heliotropium* species: (Muhaidat *et al*., 2011); *Salsola* species: (Schüssler *et al*., 2017); *Alloteropsis* species: (Lundgren *et al*., 2016; Alenazi *et al*., 2024); *Neurachninae* species: (Khoshravesh *et al*., 2020; Lauterbach *et al*., 2024)). Previous work by Khoshravesh et al. (2020) examined anatomical traits - including plasmodesmata frequency – across C_3_, C_3_-C_4_ intermediate and C_4_ species in the grass subtribe *Neurachninae*. Unlike the C_4_ crops maize and sorghum, *Neurachninae* species have a distinct Kranz anatomy (“Neurachneoid” anatomy) with two layers of BS cells, allowing the quantification of plasmodesmata between mestome sheath (MS), mesophyll (M) and parenchymatous sheath (PS)(Sedelnikova *et al*., 2018). Interestingly, C_3_-C_4_ intermediate species had intermediate plasmodesmata numbers at these interfaces, although the differences were not statistically significant.

To investigate when increases in plasmodesmata numbers arise along a more comprehensive C_3_-to-C_4_ evolutionary gradient, and in a genus that has the classical Kranz anatomy with a single BS cell layer, the dicotyledonous genus *Flaveria* is an ideal system. *Flaveria* is part of the *Asteraceae* family and includes species that use a wide range of photosynthesis types, and is a well-established model system for studying the evolution of C_4_ photosynthesis (Brown *et al*., 2005). For example, stomata frequency decreases along the C_3_-to-C_4_ gradient among *Flaveria* species, which correlates with the expression of FSTOMAGEN (Zhao *et al*., 2022). Likewise, mesophyll chloroplast numbers also decrease along this gradient (Stata *et al*., 2014, 2016). However, despite decades of work on this genus, it is not known if C_4_ *Flaveria* species have increased physical cell-to-cell connectivity between M-BS cell interfaces and if so, how this trait evolved along the C_3_-to-C_4_ evolutionary gradient. To address this gap, we applied high-resolution electron microscopy mapping, recently developed for quantifying plasmodesmata in different cell-to-cell interfaces in leaves of C_3_ and C_4_ *Cleomaceae* species (Schreier *et al*., 2024), to quantitatively assess plasmodesmata frequency in *Flaveria* species - including C_3_, C_3_-C_4_ intermediates, C_4_-like and C_4_ species. We observed a stepwise increase in plasmodesmata frequency along the C_3_ to C_4_ gradient in all cell interfaces examined, with the strongest increases in plasmodesmata numbers observed between the M-BS interface of C_4_ species, providing new insights into the evolution of this important C_4_ trait.

## Results

### Large-scale high-resolution electron microscopy provides in-depth imaging of leaf anatomy in Flaveria species

To study cell-to-cell connectivity along the C_3_-to-C_4_ gradient in *Flaveria*, we selected seven species including C_3_ (*F. cronquistii* and *F. robusta*), C_3_-C_4_ intermediate (Type I and II; *F. sonorensis* and *F. ramosissima* respectively), C_4_-like (*F. vaginata*) and C_4_ (*F. kochiana* and *F. bidentis*) species (Fig. **1A**). Based on previous phylogenetic analyses (McKown & Dengler, 2007; Lyu *et al*., 2015), the selected C_4_ and C_4_-like species and one of the C_3_-C_4_ intermediates (*F. ramosissima*) all belonged to Clade A of *Flaveria* classification, while the two C_3_ species and the other intermediate (*F. sonorensis*) were basal to this clade. Thus, the selected species likely represent an evolutionary gradient (Fig. **1A**).

**Figure 1.**
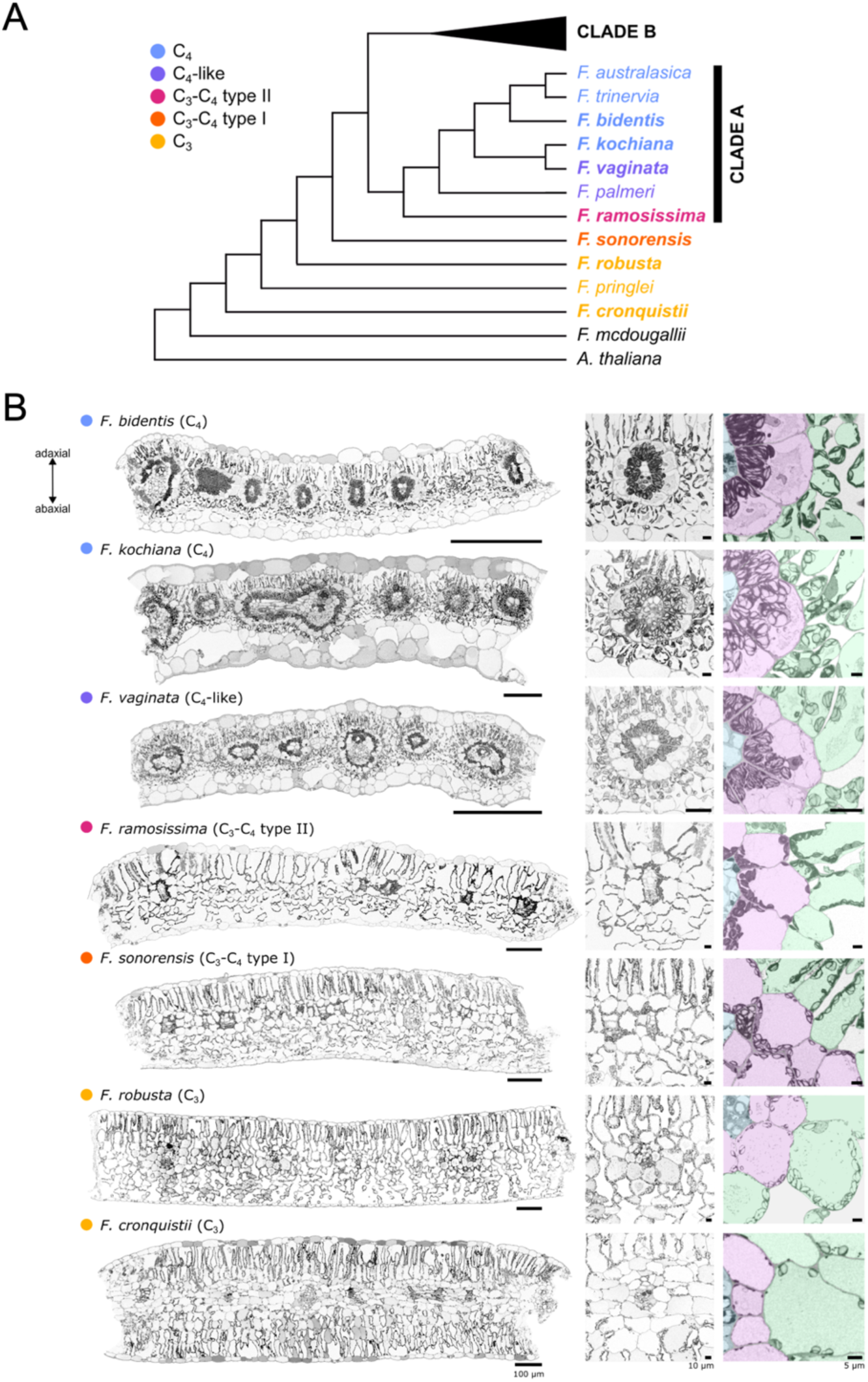
Leaf anatomy in the *Flaveria* genus along the evolutionary gradient from C_3_ to C_4_ via C_3_-C_4_ intermediate species. **(A**) Phylogenetic tree of the *Flaveria* genus based on Lyu et al (2015) and McKown et al. (2005). Different colours represent different photosynthesis types. The 7 *Flaveria* species examined in this study are in bold text. (**B**) Overview of leaf anatomy in the selected *Flaveria* species. Left panels: Representative scanning electron micrographs of mature leaf cross-sections (see Supplemental Figure S1 for mature leaves harvesting details). Cross sections are arranged so that the adaxial leaf surface is on top. Bars = 100 μm. Mid panels: Zoomed in view of a representative of an intermediate vein used for plasmodesmata quantification. Bars = 10 μm. Right panels: Zoomed in view on mesophyll (M) and bundle sheath (BS) cells depicting typical M-M, BS-BS and M-BS cell interfaces examined in this study. M cells are coloured in green, BS cells in magenta and veins in blue. Bars = 5 μm

To assess leaf anatomical traits in the seven selected *Flaveria* species, all species were grown under the same controlled environment, and fully developed, mature leaves were used for 2D scanning electron microscopy (SEM) mapping (Schreier *et al*., 2024) (Fig. **S1**). Initially, we generated maps (1000x magnification) to capture the entire cross section of the sample, covering multiple vascular bundles (Fig. **1B**; Fig. **S1**). To allow quantification, three biological replicates (with each representing a leaf from a separate plant) were imaged for each species (Fig. **S2**). The seven species displayed a wide range of leaf thickness, leaf shapes and cellular arrangements, and some typical Kranz-anatomy features were already observed in the intermediate species (Fig. **1B**; Fig. **S1**). For example, concentric M and BS cell arrangement was already visible in the intermediate *F. ramossisima*, becoming more pronounced in the C_4_-like and C_4_ species. Also, the intermediates had noticeably more chloroplasts in BS cells, which occurred more frequently towards the vein (centripetal). In the C_4_-like and C_4_ species, chloroplast placement was almost exclusively centripetal.

### Stepwise increased plasmodesmal frequency between mesophyll and bundle sheath cells from C_3_-to-C_4_ evolution

To allow visualisation and quantification of plasmodesmata, we selected areas that included intermediate vein cross sections (Fig. **S2**), and reimaged those areas at high-resolution (12,000x magnification). This approach allowed plasmodesmata to be observed in three different types of cell interface (M-BS: mesophyll – bundle sheath; M-M: mesophyll – mesophyll; BS-BS: bundle sheath – bundle sheath). Plasmodesmata frequency was quantified per unit interface length. Since plasmodesmata are typically arranged in clusters (pitfields) that vary in size (Faulkner *et al*., 2008; Danila *et al*., 2016), numbers observed can be variable depending on the section. Therefore, we assessed ∼50-150 interfaces per species for each interface type across three biological replicates. We previously demonstrated that plasmodesmal frequencies quantified this way from 2D SEM maps are comparable to those obtained from 3D electron microscopy (Schreier *et al*., 2024).

Plasmodesmata frequency at the M-BS interface was low in the C_3_ *Flaveria* species (*F. robusta* and *F. cronquistii*), and many interface sections did not contain any plasmodesmata (Fig. **2**; Table **S1**). By contrast, in C_4_-like and C_4_ species, plasmodesmata were present in almost every M-BS cell interface assessed. Both of the C_3_-C_4_ intermediate species (*F. sonorensis* and *F. ramosissima*) had significant 3-fold higher plasmodesmata frequency relative to the C_3_ species (*F. cronquistii* and *F. robusta*). There was no significant difference in plasmodesmata number between the two intermediates, which was interesting because the two C_3_-C_4_ species represent different types of intermediates (type I and II): type I relies solely on the C_2_ cycle (*F. sonorensis*) whereas type II shows some basal C_4_ cycle activity (*F. ramosissima*)(Edwards & Ku, 1987). The C_4_-like *F. vaginata* had 6-fold more plasmodesmata at the M-BS cell interface compared to C_3_ species, which was significantly greater than the C_3_-C_4_ intermediate species. The C_4_ species, *F. bidentis* and *F. kochiana*, showed 9-fold and 7-fold higher plasmodesmata numbers, respectively, compared to C_3_ species. Therefore, our quantification revealed that plasmodesmata numbers at the M-BS interface increased stepwise along the C_3_-to-C_4_ evolutionary gradient in our selected *Flaveria* species

**Figure 2.**
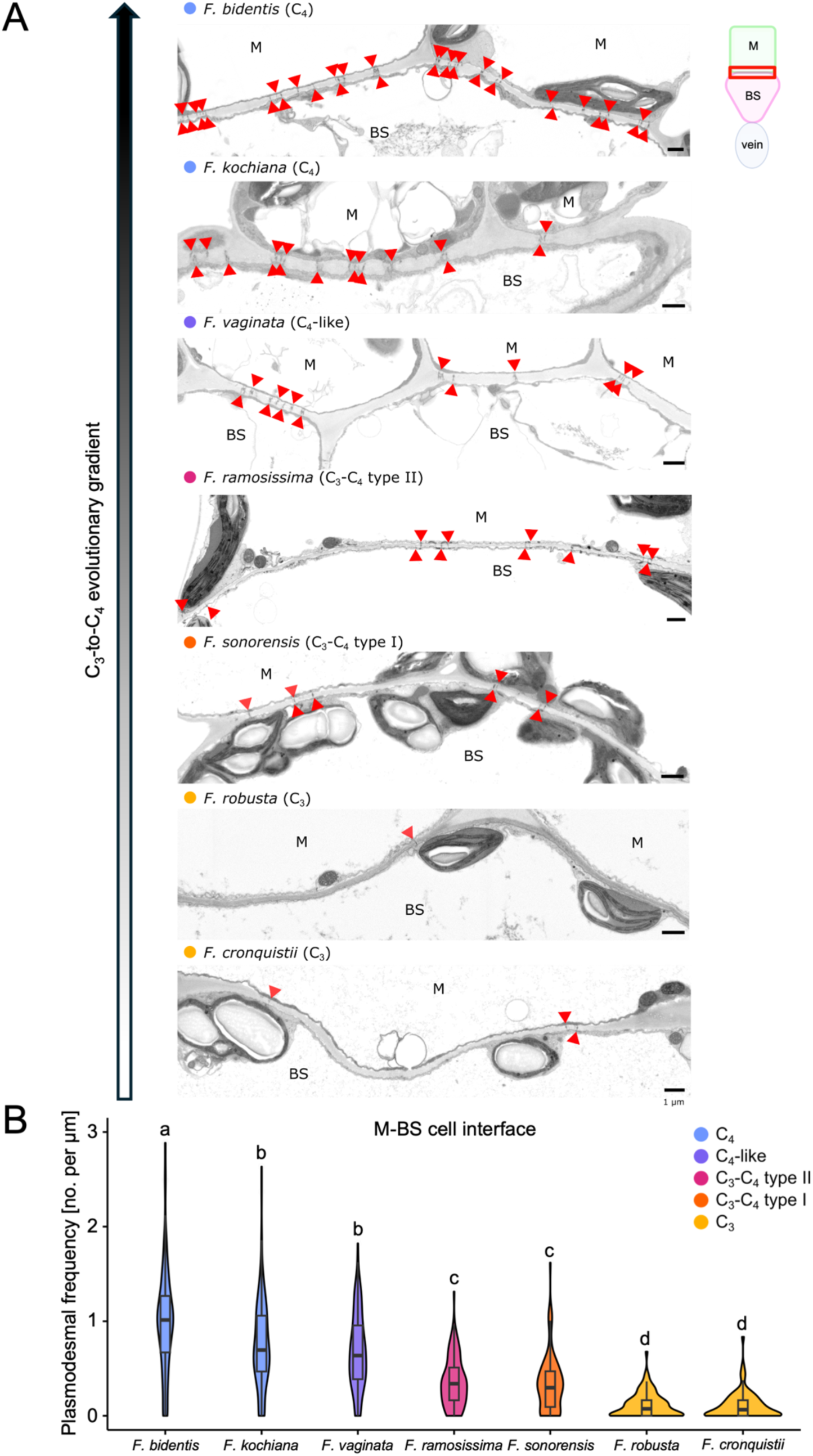
Increase in plasmodesmata number at the mesophyll (M) – bundle sheath (BS) cell interface along the C_3_-to-C_4_ evolutionary gradient in *Flaveria* genus. **(A)** Representative excerpts from 2D scanning electron microscopy map, showing M and BS cell interfaces in all seven *Flaveria* species. Excerpts are oriented that M cells are on the top and BS cells are on the bottom, as shown in the diagram on the right. Red arrows mark individual plasmodesmata. Bars = 1 μm (**B**) Violin plot of plasmodesmal frequency quantified at the M-BS cell interface in mature leaf samples. BS surrounding intermediate veins were chosen for plasmodesmata quantifications. Plasmodesmal frequencies were quantified in 79-156 individual M-BS cell interfaces per species, across three individual biological replicates. Box and whiskers represent the 25 to 75 percentile and minimum-maximum distributions of the data. Letters show statistical ranking using a Kruskal-Wallis test, followed by Dunn’s post hoc test (with different letters indicating statistically differences of *P<0.01*). Values indicated by the same letter are not statistically different.

Given the strong increase in plasmodesmata frequency at the M-BS interface along the C_3_-to-C_4_ evolutionary gradient, we examined plasmodesmata numbers at the M-M and BS-BS interfaces. At the BS-BS interface, the C_4_ species *F. bidentis* had a significant increase in plasmodesmata numbers compared to the C_3_ species, but the increase was less than 2-fold (Fig. **3A, C**; Table **S1**). Unfortunately, plasmodesmata could not be quantified in C_4_ *F. kochiana*, because the BS-BS cell interface was too electron-dense compared to other interfaces, making it difficult to distinguish individual plasmodesmata (Fig. **S3**). The reason for this is not known but may reflect a difference in cell wall composition or properties. At the M-M interfaces, plasmodesmal frequency increased 3-fold from C_3_ species to C_3_-C_4_ intermediate species, 4-fold to C_4_-like species, and 5 to 6-fold in the C_4_ species *F. kochiana* and C_4_ *F. bidentis*, respectively (Fig. **3B,D**; Table **S1**). Therefore, the stepwise increase in plasmodesmata frequency along the C_3_-to-C_4_ evolutionary gradient was also apparent at the M-M interface, but not at the BS-BS interface.

**Figure 3.**
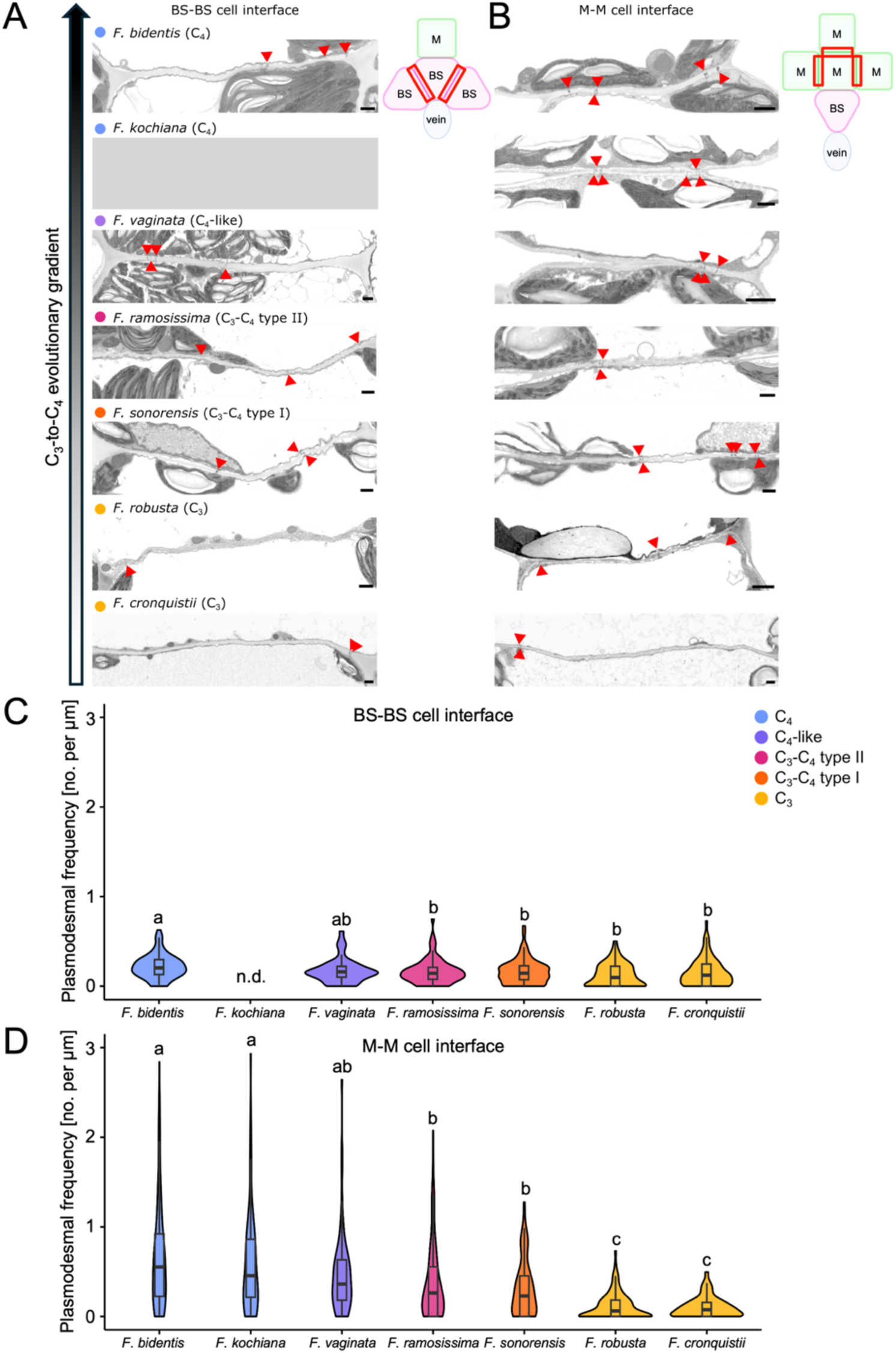
Plasmodesmal frequency in bundle sheath – bundle sheath (BS-BS) and mesophyll - mesophyll (M-M) cell interfaces along the C_3_-to-C_4_ evolutionary gradient in *Flaveria* species. **(A)** Representative excerpts from 2D scanning electron microscopy maps showing BS-BS cell interfaces in six of seven *Flaveria* species. An illustration is provided to show the position of the interface. Red arrows indicate individual plasmodesmata. Bars = 1 μm. **(B)** As for A, except for M-M cell interfaces in all seven *Flaveria* species. (**C**) Violin plot of plasmodesmal frequency measured at BS-BS cell interface in mature leaf samples. Plasmodesmal frequencies were quantified from a range of 54-84 individual BS-BS cell interfaces per species across three individual biological replicates per species. Box and whiskers represent the 25 to 75 percentile and minimum-maximum distributions of the data. Letters show statistical ranking using a Kruskal-Wallis test, followed by Dunn’s post hoc test (with different letters indicating statistically differences of *P<0.05*). Values indicated by the same letter are not statistically different. n.d. = not determined (**D**) As for C, but for M-M cell interfaces. Plasmodesmal frequencies were quantified from 68-116 individual M-M cell interfaces per species across three biological replicates.

Interestingly, plasmodesmata frequency was equal at the M-M and M-BS interfaces in all C_3_ and C_3_-C_4_ intermediate species. Thus, C_3_-C_4_ intermediates can be characterised as having a general increase in cell-to-cell connectivity relative to C_3_ species, which may be sufficient for facilitating C_2_ metabolism. This feature is distinct from the polar plasmodesmata enrichment observed in C_4_ and C_4_-like species, where plasmodesmata frequency was always greater at the M-BS interface compared to the M-M interface.

### Bundle sheath chloroplast coverage correlates with plasmodesmal frequency along C_3_-to-C_4_ evolutionary trajectory

We assessed whether the observed increase in plasmodesmata correlates with other BS cell characteristics along the C_3_-to-C_4_ gradient (Fig. **4**; Fig. **S4**; Table **S2**). We quantified BS cell area (Fig. **4A,B**), number of chloroplasts per BS cell (Fig. **4A,C**), chloroplast area per BS cell (Fig. **4A,D**), and total BS chloroplast coverage (Fig. **4A,E**). There was no clear trend in BS cell area along the gradient. The number of chloroplasts per bundle sheath cell were sharply higher in the C_3_-C_4_ intermediate, C_4_-like and C_4_ species compared to the two C_3_ species, but there were no differences in the number of chloroplasts between the C_3_-C_4_ intermediate and C_4_ species. However, when we quantified the total chloroplast area within a bundle sheath cell (BS chloroplast coverage), we observed a stepwise increase along the C_3_-to-C_4_ evolutionary gradient, as the C_4_ species had larger chloroplasts (Fig. **4A**; Fig. **S4A**, **S5**). Thus, the higher cell-to-cell connectivity at the M-BS interfaces coincides with the increases in chloroplast area within BS cells, which could be seen in the high correlation coefficients between chloroplast coverage and plasmodesmata numbers (Fig. **S6**).

**Figure 4.**
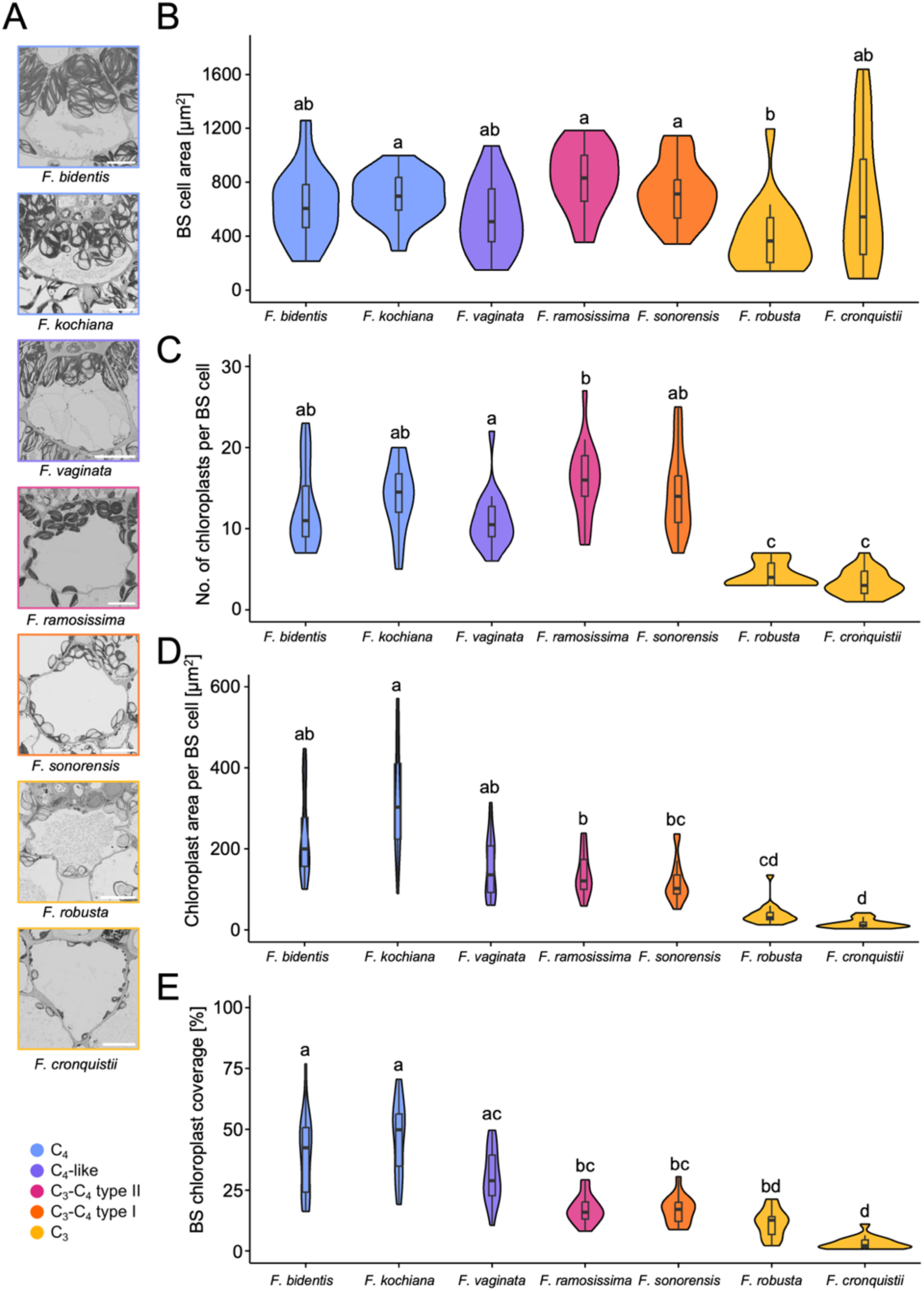
Bundle sheath (BS) anatomy along C_3_-to-C_4_ evolutionary gradient in *Flaveria*. **(A**) Scanning electron micrograph excerpts of representative BS cells in all seven *Flaveria* species. Bars = 10 μm (**B**) Violin plot of BS cell area in the different *Flaveria* species. BS cell areas were quantified from 18 individual BS cells per species (6 BS per biological replicate). Box and whiskers represent the 25 to 75 percentile and minimum-maximum distributions of the data. Letters show statistical ranking using a Kruskal-Wallis test, followed by Dunn’s post hoc test (with different letters indicating statistically differences of *P<0.05*). Values indicated by the same letter are not statistically different. (**C**) Violin plot of number of chloroplasts per BS cell in each *Flaveria* species. Letters show statistical ranking using an one-way ANOVA with a Tukey’s post hoc test (with different letters indicating statistically differences of *P<0.05*). (**D**) Violin plot of chloroplast area per BS cell area covered by chloroplasts (BS chloroplast coverage) in percentage for each *Flaveria* species. Letters show statistical ranking using a Kruskal-Wallis test, followed by Dunn’s post hoc test (with different letters indicating statistically differences of *P<0.05*). (**E**) Violin plot of BS chloroplast coverage across different *Flaveria* species. The value was quantified from 18 individual BS cells per species (6 BS per biological replicate). Letters show statistical ranking using a Kruskal-Wallis test, followed by Dunn’s post hoc test (with different letters indicating statistically differences of *P<0.05*).

## Discussion

### Enhanced plasmodesmal frequency is a conserved C_4_ trait and evolved stepwise in Flaveria genus

Using the *Flaveria* genus, we observed a stepwise increase in plasmodesmata frequency along a C_3_-to-C_4_ gradient. The increase in plasmodesmata at the M-BS interface is a common trait among different C_4_ lineages, although there is variation in the degree of increase between species, which may depend on C_4_ subtype (Danila *et al*., 2016, 2018; Schreier *et al*., 2024). C_4_ *Flaveria* species conduct the NADP-ME subtype of C_4_ photosynthesis, and our plasmodesmata quantification method revealed a 7 to 9-fold higher plasmodesmal frequency at the M-BS interface in the C_4_ species (*F. kochiana* and *F. bidentis*) compared to the C_3_ species (Fig. **2**; Table **S1**). This increase is comparable to the 6-to 9-fold higher plasmodesmata numbers compared to C_3_ grasses reported for C_4_ grasses conducting the NADP-ME C_4_ subtype (Botha, 1992; Danila *et al*., 2016, 2018). However, species conducting the NAD-ME C_4_ subtype have the most drastic increase in plasmodesmata numbers at M-BS cell interfaces compared to the NADP-ME and PCK subtypes (Danila *et al*., 2018). For example, we previously observed up to 13-fold higher plasmodesmata number in *G. gynandra* (NAD-ME subtype) compared to its close C_3_ relatives (Schreier *et al*., 2024). Plasmodesmal frequencies in our C_3_ species (*F. cronquistii* and *F. robusta*) were comparable to values reported for other C_3_ species, *Arabidopsis thaliana* and *Tarenaya hassleriana* (Fig. **2,3**; Table **S1**; (Schreier *et al*., 2024)). Interestingly, *F. robusta* was reported to have some proto-Kranz anatomy features such as increased BS organelle development (Fig. **4A,C****,D**;(Sage *et al*., 2013)). The comparable plasmodesmal frequency in this species compared to *F. cronquistii* and other distantly related C_3_ eudicots suggests that the degree of proto-Kranz anatomy in *F. robusta* does not include increased leaf plasmodesmal frequencies.

In the C_3_-C_4_ intermediate species (*F. sonorensis* and *F. ramosissima*), plasmodesmal frequency was significantly increased in both M-BS and M-M interfaces compared to C_3_ species, leading to similar plasmodesma numbers between the two interfaces (Fig. **2,3**; Table **S1**). It was only in the C_4_-like species *F. vaginata* and the two C_4_ species (*F. kochiana* and *F. bidentis*) where a strong increase in plasmodesmata numbers was observed in the M-BS interface compared to the M-M cell interface. This suggests that the polar formation of plasmodesmata that leads to the very high plasmodesmata frequency specifically at the M-BS cell interface in C_4_-like/C_4_ species, occurs as a second evolutionary step from general increase in plasmodesmata frequency observed in the C_3_-C_4_ intermediate species, and coincides with the evolution of C_4_ metabolism. The stepwise increase in the *Flaveria* genus could thus be exploited as a model system to identify factors specifically involved in polar plasmodesmata formation at the M-BS interface, for example, by comparing the molecular players in plasmodesmata formation between C_3_-C_4_ intermediate and C_4_-like/C_4_ species.

### Cell-to-cell connectivity and metabolite shuttling in C_3_-to-C_4_ evolution

The increase in physical cell-to-cell connectivity between M and BS cells along the C_3_-to-C_4_ evolutionary gradient coincides with the photosynthetic pathway becoming more reliant on efficient metabolite shuttling between these two cell types. Metabolite profiling of *Flaveria* species found multiple metabolite shuttles operating between M and BS cells (Borghi *et al*., 2022). The NADP-ME C_4_-photosynthesis pathway operates in the C_4_-like and C_4_ *Flaveria* species, and to a much lesser extent in the C_3_-C_4_ intermediate type II *F. ramosissima* (Drincovich *et al*., 1998), thereby shuttling malate/alanine across the two cell types to build a carbon concentrating mechanism (Borghi *et al*., 2022)(Fig. **5**). Intermediate C_3_-C_4_ species conduct a “C_2_-type” of photosynthesis, where part of the photorespiratory pathway (glycine decarboxylase complex) is exclusively located in the BS cells, leading to a modest carbon concentrating mechanism around RuBisCO in BS cells (Keerberg *et al*., 2014; Lundgren, 2020; Mercado & Studer, 2022). There is much diversity in the how the C_2_ photosynthesis pathway operates among the independent C_3_-C_4_ evolutionary lineages. In C_3_-C_4_ intermediate species that conduct C_2_-type photosynthesis, glycine and serine are shuttled between M and BS cells (Borghi *et al*., 2022). Additionally, aspartate may be shuttled in the C_3_-C_4_ intermediate species *F. ramosissima* and C_4_-like species *F. palmeri* (Borghi *et al*., 2022). Therefore, the carbon-concentrating mechanism of C_3_-C_4_ intermediate species conducting C_2_ photosynthesis also relies on cell-to-cell connectivity between M and BS cells.

**Figure 5.**
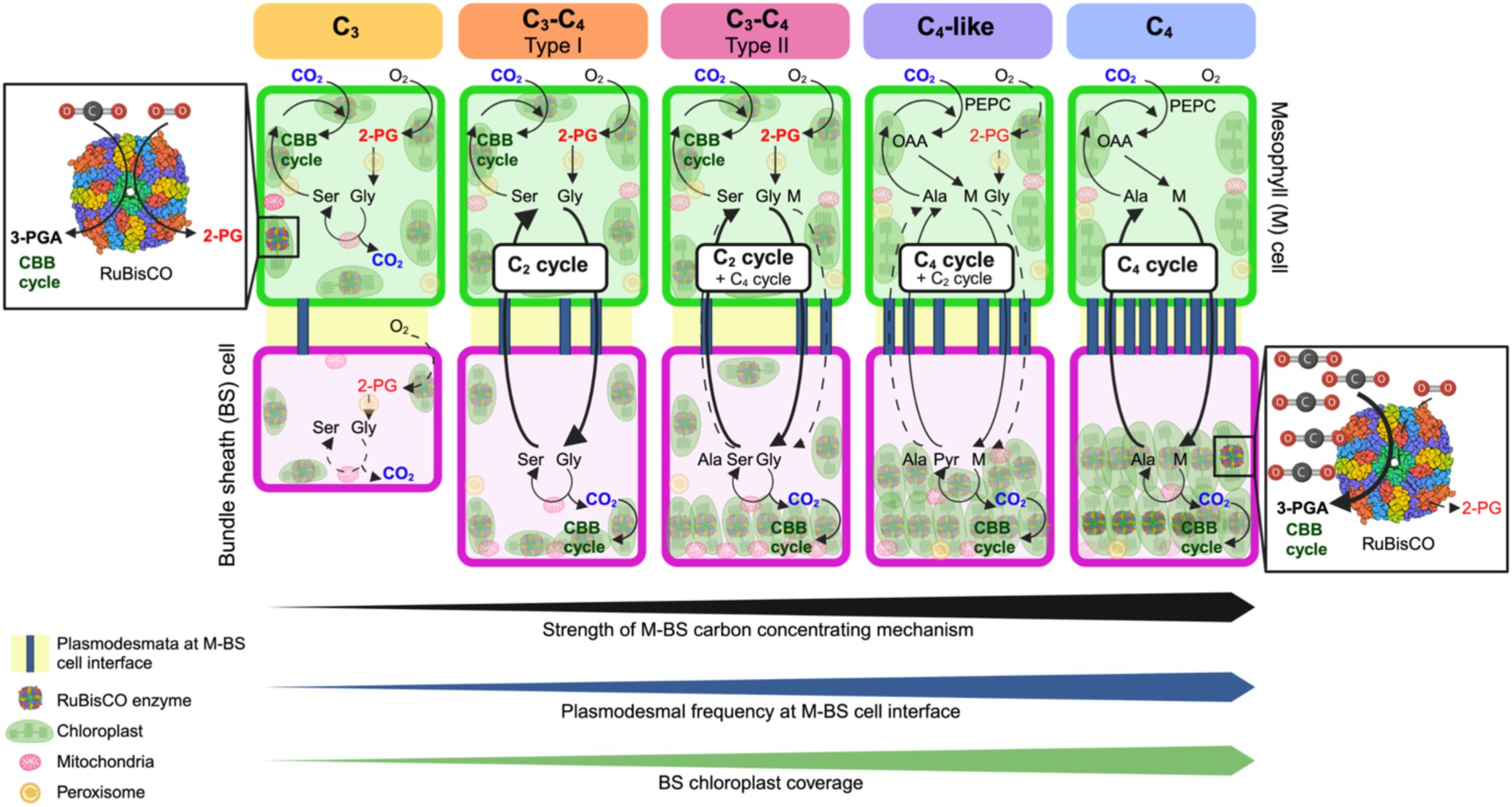
A model of increasing cell-to-cell connectivity along the C_3_-to-C_4_ evolutionary gradient via C_3_-C_4_ intermediates to accommodate metabolite shuttles between mesophyll (M, green) and bundle sheath (BS, magenta) cells. C_3_-C_4_ intermediate species run a C_2_ cycle, as part of the photorespiration pathway (glycine decarboxylation complex) is moved exclusively to the BS cells, thereby shuttling C_2_ acid (glycine - Gly) and serine (Ser) between M and BS cells. C_4_-like and C_4_ species run a NADP-ME type of C_4_ cycle, where C_4_ acid (malate - M) and C_3_ acid (pyruvate - Pyr) are shuttled between M and BS cells to concentrate CO_2_ around RuBisCO. RuBisCO: Ribulose-1,5-Bisphophate Carboxylase/Oxygenase; CBB cycle: Calvin-Benson-Bassham cycle; 3-PGA: 3-phosphoglyceric acid; 2-PG: 2-Phosphoglycolate; Ser: Serine; Gly: Glycolate; OAA: Oxaloacetate; M: Malate; Ala: Alanine.

Consequently, the increase in cell-to-cell connectivity, as evidenced by the rise in plasmodesmal frequency, likely represents a structural adaptation to meet the metabolic demands of these photosynthesis pathways (Fig. **5**). Further research is needed to understand whether increased cell-to-cell connectivity was a prerequisite for the development of C_2_ and C_4_ carbon concentrating mechanisms.

### Metabolic link between chloroplasts and plasmodesmata formation

In addition to enhanced connectivity between M and BS cells, Kranz anatomy is also characterised by increased organelle development in BS cells, reflecting the important metabolic role in C_4_ photosynthesis (Sedelnikova *et al*., 2018). The increase in plasmodesmal numbers at the M-BS interface along the C_3_-to-C_4_ gradient coincides with the increase in chloroplast coverage in bundle sheath cells (Fig. **2,4**; Table **S1,S2**). The chloroplast quantification values that we obtained for BS cells are comparable to those reported by Stata *et al*. (2014, 2016).

This observation is interesting because there is genetic evidence for a link between organelle and plasmodesmata development. A key transcription factor involved in chloroplast development in maize (GOLDEN2) was shown to promote both organelle development and plasmodesmata numbers when expressed in rice (Wang *et al*., 2017). Recent findings also indicate that plasmodesmata are regulated by signals originating from organelles, mainly chloroplasts and mitochondria. This pathway, named <u>o</u>rganelles-<u>n</u>ucleus-<u>p</u>lasmodesmata-<u>s</u>ignalling (ONPS), acts on the nucleus to modulate the expression of genetic pathways that control both plasmodesmata regulation and formation (Ganusova *et al*., 2020; Azim & Burch-Smith, 2020). Possible examples of the signals include the maize gene *SXD1*, which encodes a chloroplast-localized tocopherol cyclase (VTE1 homolog) that is associated with callose deposition around plasmodesmata at the bundle sheath and vascular parenchyma cell interface (Botha *et al*., 2000; Provencher *et al*., 2001). In Arabidopsis, the increased size exclusion limit mutants (*ise1* and *ise2*), which show increased cell-to-cell transport at the mid-torpedo stage of embryogenesis, are also both defective in organellar proteins (Kobayashi *et al*., 2007; Stonebloom *et al*., 2009). ISE1 encodes a mitochondrial-localised DEAD-box RNA helicase (Stonebloom *et al*., 2009) and ISE2 a chloroplast-localised RNA helicase (Kobayashi *et al*., 2007; Burch-Smith & Zambryski, 2010). Furthermore, GFP arrested trafficking (GAT1) is a plastidial thioredoxin, which when mutated restricts GFP transport from the phloem to the root meristem by increasing callose deposition (Benitez-Alfonso *et al*., 2009).

Importantly, we recently showed that plasmodesmata are formed at the M-BS cell interface after light induction in dark-grown cotyledons of C_4_ *G. gynandra* seedlings. De-etiolation represents a transition from heterotrophic to autotrophic metabolism, therefore relying on active photosynthesis, and triggering a rapid increase in plasmodesmata formation. In this system, chemical inhibition of chloroplast development and photosynthesis prevented the increase in plasmodesmata numbers at the M-BS cell interface, suggesting that functional chloroplasts and active photosynthesis is important for their formation (Schreier *et al*., 2024). Thus, plasmodesmata formation in C_4_ *G. gynandra* is linked to the induction of C_4_ metabolism and thus shuttling of C_4_/C_3_ acids between M and BS cells. It is tempting to speculate that this light-induced mechanism of increasing plasmodesmata at the M-BS cell interface evolved along the C_3_-to-C_4_ evolutionary gradient and may underlie the second step in the observed increase in plasmodesmata numbers between C_3_-C_4_ intermediate species and C_4_-like/C_4_ Flaveria species. The molecular mechanism that links plasmodesmata formation and chloroplast development in BS cells requires further investigation.

## Material and methods

### Plant material and growth conditions

All *Flaveria* species were grown were on soil in pots in a walk-in climate-controlled growth chamber at the plant growth facility at the Department of Plant Sciences (Cambridge). The chamber was set to 16 h/8 h light/dark cycles, with 350 μmol photons m^−2^ s^−1^ light intensity, constant temperature of 22°C, 65% (v/v) relative humidity, and ambient CO_2_. *Flaveria* species were propagated from cuttings: Healthy plants with shoots containing at least two nodes or sets of matured leaves was selected. Shoots were cut using sharp scissors or a razor blade, and lower leaves were removed. The remaining leaves were trimmed to half their size, then a diagonal cut at the bottom of the stems were made. The cut end was dipped in rooting powder (Karrma Ltd) and the cuttings were placed in pre-moistened soil. The potted cuttings were covered by a lid with closed vents for two weeks or until new growth appeared, then the vents were gradually opened to allow further growth.

### Sample preparation for electron microscopy

To assess the leaf anatomy, samples from 3-6 individual plants for each *Flaveria* species were harvested for electron microscopy. All samples were harvested towards the end of the photoperiod (∼12 h into the 16 h photoperiod). A leaf segment (∼2 mm^2^) was excised using a razor blade from the midpoint of the proximal-distal axis of the leaf and between the midrib and the leaf edge. One sample was harvested per individual plant and immediately fixed in 2% (v/v) glutaraldehyde and 2% (w/v) formaldehyde in 0.05 - 0.1 M sodium cacodylate (NaCac) buffer (pH 7.4) containing 2 mM calcium chloride. After 12-24 h of fixation at 4°C, fixed samples were washed 5 times in 0.05 – 0.1 M NaCac buffer, and post-fixed in 1% (v/v) aqueous osmium tetroxide, 1.5% (w/v) potassium ferricyanide in 0.05 M NaCac buffer for 3 days at 4°C. After osmication, samples were washed 5 times in deionized water and post-fixed in 0.1% (w/v) thiocarbohydrazide for 20 min at room temperature in the dark. Samples were then washed 5 times in deionized water and osmicated for a second time for 1 h in 2% (v/v) aqueous osmium tetroxide at room temperature. Samples were washed 5 times in deionized water and subsequently stained in 2% (w/v) uranyl acetate in 0.05 M maleate buffer (pH 5.5) for 3 days at 4°C, and washed 5 times afterwards in deionized water. Samples were then dehydrated in an ethanol series, transferred to acetone, and then to acetonitrile. Leaf samples were embedded in Quetol 651 resin mix (TAAB Laboratories Equipment Ltd) and cured at 60°C for 2 days.

### Scanning electron microscopy (SEM)

For 2D SEM mapping, ultra-thin sections were placed on Melinex (TAAB Laboratories Equipment Ltd) plastic coverslips mounted on aluminium SEM stubs using conductive carbon tabs (TAAB Laboratories Equipment Ltd), sputter-coated with a thin layer of carbon (∼ 30 nm) to avoid charging and imaged in a Verios 460 scanning electron microscope at 4 keV accelerating voltage and 0.2 nA probe current using the concentric backscatter detector in field-free (low magnification) or immersion (high magnification) mode (working distance 3.5 – 4 mm, dwell time 3 µs, 1536 × 1024 pixel resolution). For plasmodesmata frequency quantification, SEM stitched maps were acquired at 12,000X magnification using the FEI MAPS automated acquisition software. Greyscale contrast of the images was inverted to allow easier visualisation.

### Plasmodesmata quantification

Plasmodesmal frequency from 2D electron microscopy maps was determined as previously described (Schreier *et al*., 2024). Briefly, plasmodesmal frequency was determined as the number of plasmodesmata observed per μm length of shared cell interface between two cell types (mesophyll – bundle sheath, mesophyll – mesophyll, bundle sheath – bundle sheath). Plasmodesmata numbers and cell lengths were determined using ImageJ software. Plasmodesmata were defined as dark channels in the EM images. Depending on plasmodesmata orientation, the entire channel was sometimes not visible on 2D EM images, and so only channels that spanned more than half of the cell wall width were counted. Branching structures of plasmodesmata (in the case of Y-shaped, branched/twinned and complex plasmodesmata) were counted as single plasmodesmata if connected.

### Chloroplast quantification

Adobe Photoshop for iPad software (version 6.0.3) was used to perform segmentation by manually marking regions of interest (i.e. bundle sheath cell area, mesophyll cell area, bundle sheath chloroplast area). Marked images were transferred to Fiji (version 1.54p) and underwent thresholding to produce a binary image from which areas (µm^2^) were extracted. Six cells per sample were randomly selected for bundle sheath total cell area and the associated chloroplast area measurements.

Chloroplast coverage was calculated as:

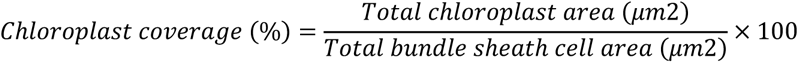

Average BS chloroplast size was calculated as:

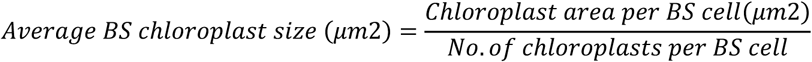

All cells were marked and quantified to determine the bundle sheath-mesophyll cell ratio. Bundle sheath cells were identified as those in closest proximity to the vascular cylinder, while mesophyll cells were identified based on size, shape and chloroplast characteristics.

### Statistical analysis

Quantification datasets were tested for normality (Shapiro-Wilk test, p > 0.05) and homogeneity of variance (Barlett’s test, p > 0.05). Normally distributed datasets were analysed using a one-way analysis of variance (ANOVA), followed by Tukey’s post hoc test. Non-normally distributed data were analysed using the Kruskal-Wallis test, followed by Dunn’s post hoc test. All statistical analyses were conducted in RStudio (version 2023.12.1+402).

## Competing interests

None declared.

## Authors contributions

TBS conceived, directed the research, and performed the experiments. TBS and KA performed the image analysis. TBS wrote the article with inputs from KA.

## Acknowledgments

The work was funded by the BBSRC Discovery Fellowship (grant CBR00550 to T.B.S.). We thank Dr. Tianshu Sun (University of Cambridge) for growing the *Flaveria* species and Prof. Julian Hibberd (University of Cambridge) for providing the growth space. We thank Dr. Karin H. Müller, Georgina E. Lindop, Dr. Filomena Gallo, and Dr. Melissa J. Drignon from the Cambridge Advanced Imaging Centre (CAIC) for the electron microscopy sample preparation as well as the support during the SEM mapping image acquisition. I also thank Prof. Julian Hibberd, Prof. Jane Langdale, Dr. David Seung and all members of the Langdale group for constructive discussions of the results.

---

**Supplemental Table S1.**
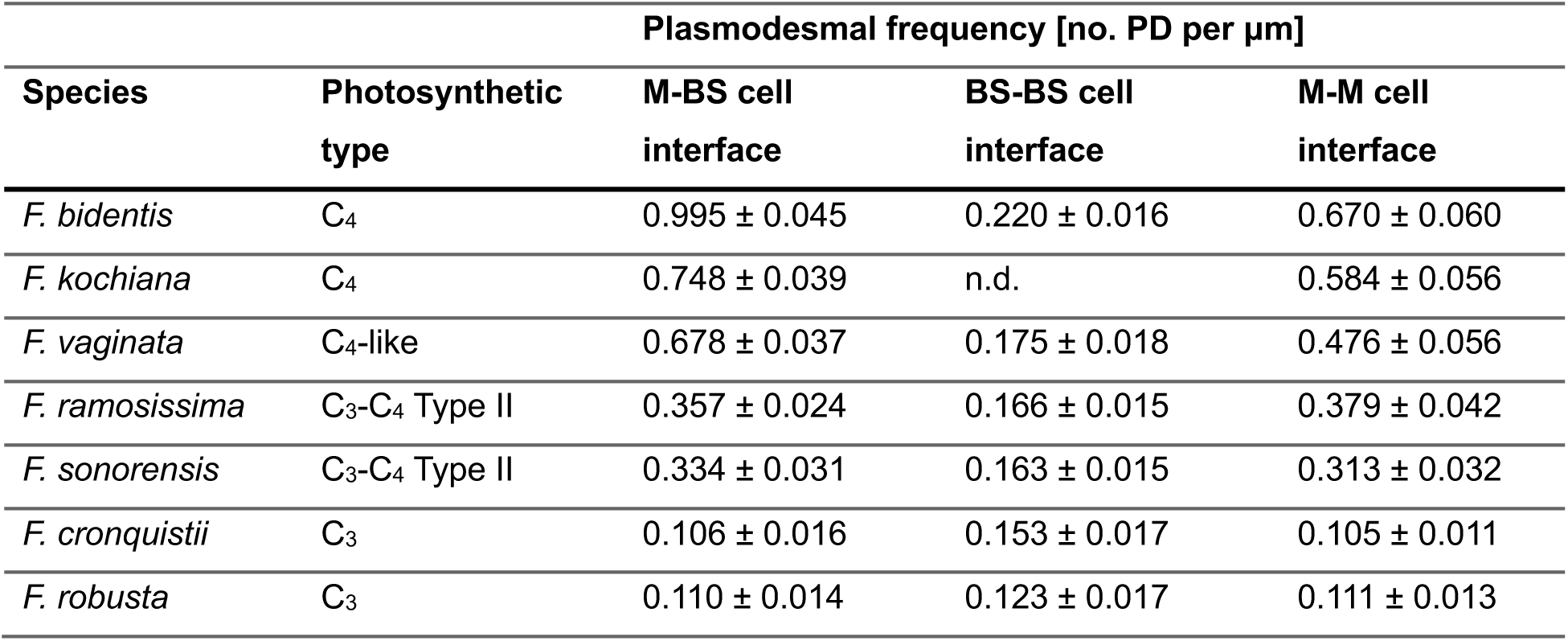
Plasmodesmal frequency in seven *Flaveria* species, measured at mesophyll-bundle sheath (M-BS), mesophyll-mesophyll (M-M), and bundle sheath-bundle sheath (BS-BS) cell interfaces. Mean values ± standard deviation of mean (standard error, SE). n.d. = not determined

**Supplemental Table S2.** Bundle sheath (BS) chloroplast quantification in seven *Flaveria* species. Mean values ± standard deviation of mean (standard error, SE).

| Species | Photosynthetic type | BS cell area [$\mu\text{m}^2$] | Chloroplast area per BS cell [$\mu\text{m}^2$] | BS chloroplast coverage [%] | No. of chloroplasts per BS cell | Average chloroplast size [$\mu\text{m}^2$] |
| --- | --- | --- | --- | --- | --- | --- |
| *F. bidentis* | C<sub>4</sub> | 633.67 ± 62.20 | 230.71 ± 24.97 | 39.66 ± 3.82 | 12.44 ± 1.18 | 18.50 ± 0.91 |
| *F. kochiana* | C<sub>4</sub> | 696.42 ± 41.43 | 324.41 ± 32.34 | 46.67 ± 3.60 | 13.83 ± 0.99 | 23.28 ± 1.33 |
| *F. vaginata* | C<sub>4</sub>-like | 544.51 ± 60.89 | 154.65 ± 17.85 | 30.59 ± 2.57 | 11.00 ± 0.87 | 13.66 ± 0.94 |
| *F. ramosissima* | C<sub>3</sub>-C<sub>4</sub> Type II | 836.74 ± 58.20 | 138.21 ± 12.74 | 17.01 ± 1.36 | 15.94 ± 1.12 | 8.62 ± 0.43 |
| *F. sonorensis* | C<sub>3</sub>-C<sub>4</sub> Type II | 723.59 ± 56.56 | 119.29 ± 13.28 | 16.82 ± 1.43 | 14.38 ± 1.28 | 8.38 ± 0.55 |
| *F. robusta* | C<sub>3</sub> | 394.26 ± 60.82 | 38.56 ± 6.50 | 11.47 ± 1.33 | 4.33 ± 0.37 | 9.02 ± 1.02 |
| *F. cronquistii* | C<sub>3</sub> | 665.18 ± 114.94 | 15.61 ± 2.71 | 3.21 ± 0.61 | 3.28 ± 0.38 | 4.44 ± 0.48 |

**Supplemental Figure S1.**
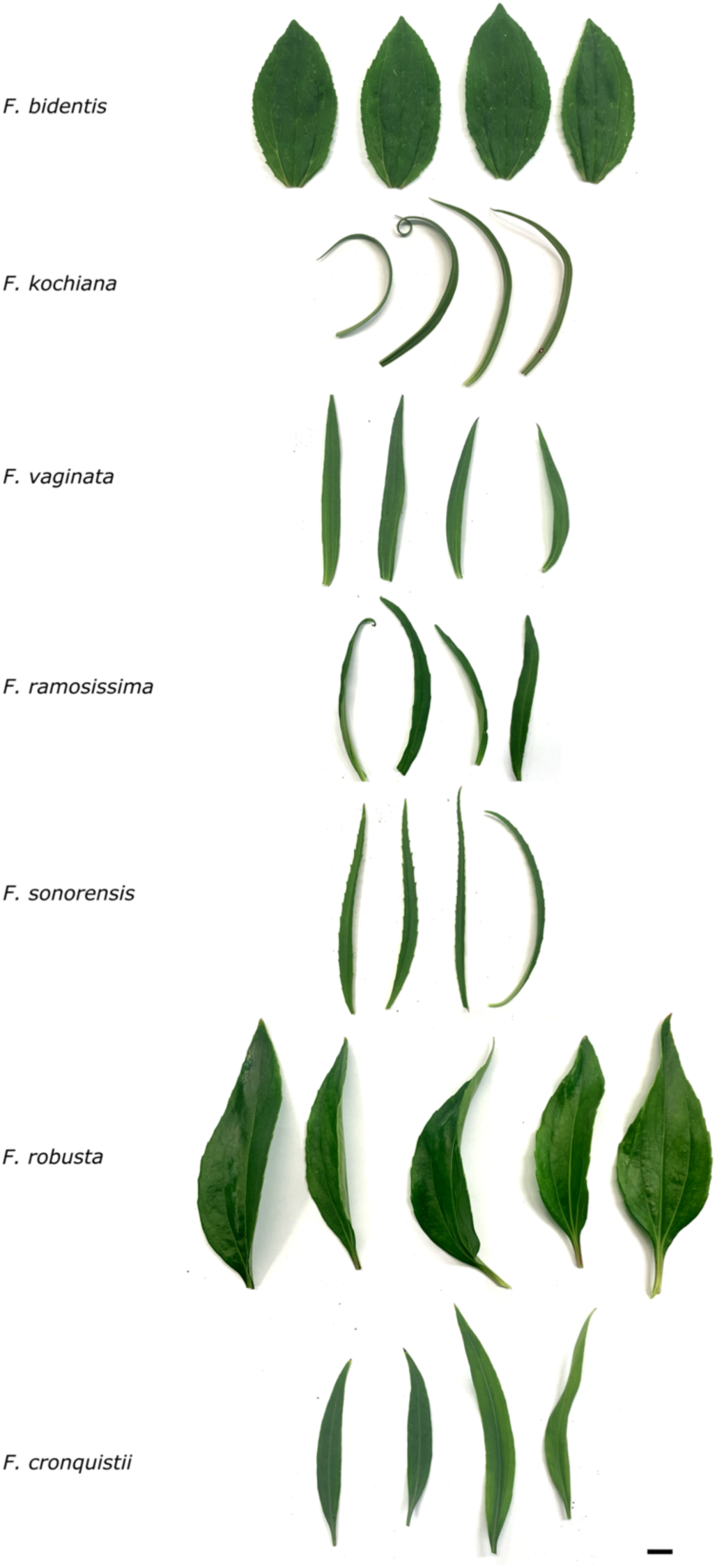
Photographs of mature leaves harvested from *Flaveria* species. Bar = 1 cm

**Supplemental Figure S2:**
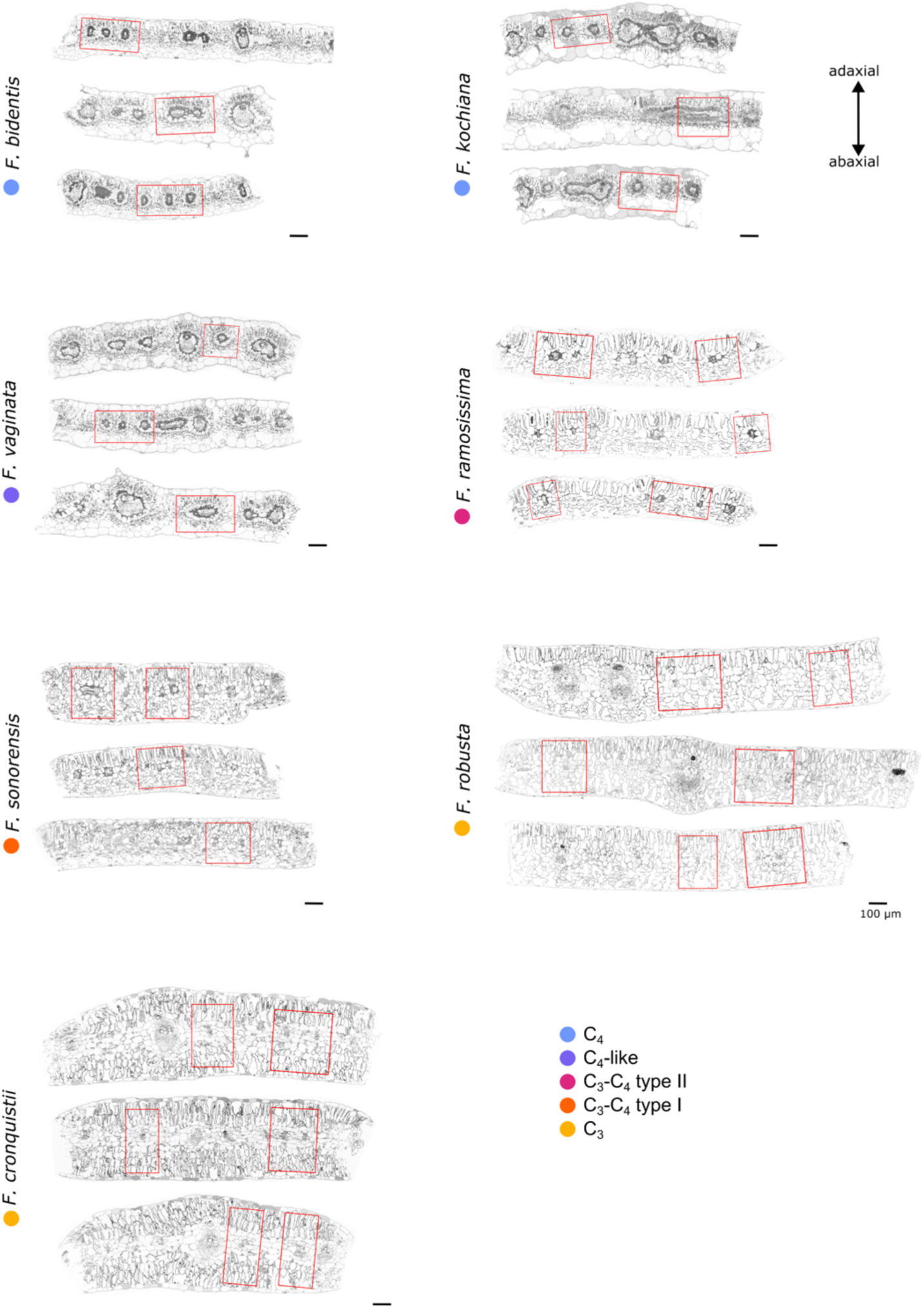
Overview of leaf anatomy from three biological replicates per *Flaveria* species. Scanning electron micrographs (1000x magnification) of mature leaf cross-sections are shown (see Supplemental Figure S1 for details on mature leaves harvested and imaged). Cross sections are oriented with the adaxial leaf surface at the top. Areas around vein cross sections used for high-resolution SEM mapping are indicted with red boxes. Bars = 100 μm

**Supplemental Figure S3:**
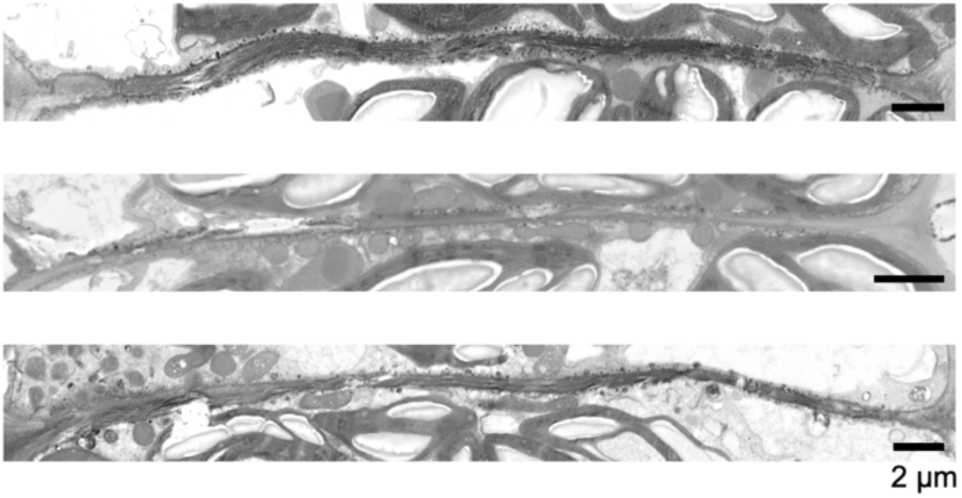
Representative excerpts from 2D scanning electron microscopy maps – one from each biological replicate - showing electron-dense bundle sheath-bundle sheath (BS-BS) cell interfaces in *F. kochiana.* No plasmodesmata were quantified at this leaf cell interface for this *Flaveria* species. Bars = 2 μm

**Supplemental Figure S4.**
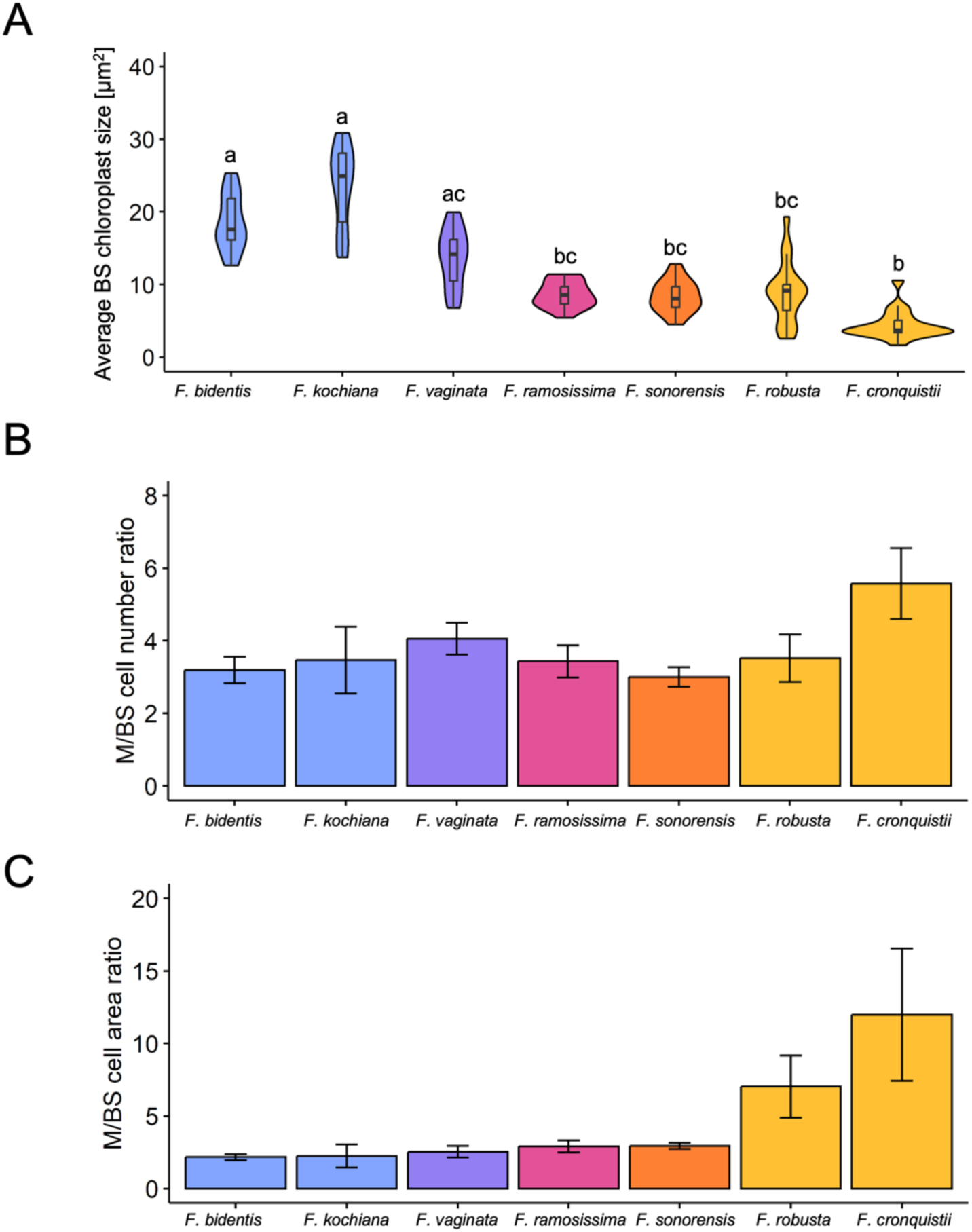
Additional quantified bundle sheath (BS) cell and Kranz anatomy traits along C_3_-to-C_4_ evolutionary gradient in *Flaveria*. **(A**) Violin plot of average BS chloroplast size across the different *Flaveria* species. These values were calculated as follows: Total chloroplast area per BS cell [μm^2^] divided by the number of chloroplasts per BS cell. Average BS chloroplast size was quantified from 18 individual BS cells per species (6 BS per biological replicate). Box and whiskers represent the 25 to 75 percentile and minimum-maximum distributions of the data. Letters show statistical ranking using an one-way ANOVA with a Tukey’s post hoc test (with different letters indicating statistically differences of *P<0.05*). Values indicated by the same letter are not statistically different. (**B**) The bar plot represents the combined M/BS cell number ratio from the three biological replicates per species. For each biological replicate, M/BS cell number was quantified from the area of one high-resolution scanning electron microscopy (SEM) map per leaf cross section. The bar plot represents the combined M/BS cell number ration from the three biological replicates per species. No statistically significant difference (*P<0.05)* were found between species using an one-way ANOVA. (**C**) Same as (B), but showing the ratio of M to BS cell area. No statistically significant difference (*P<0.05)* were found between species using an one-way ANOVA.

**Supplemental Figure 5:**
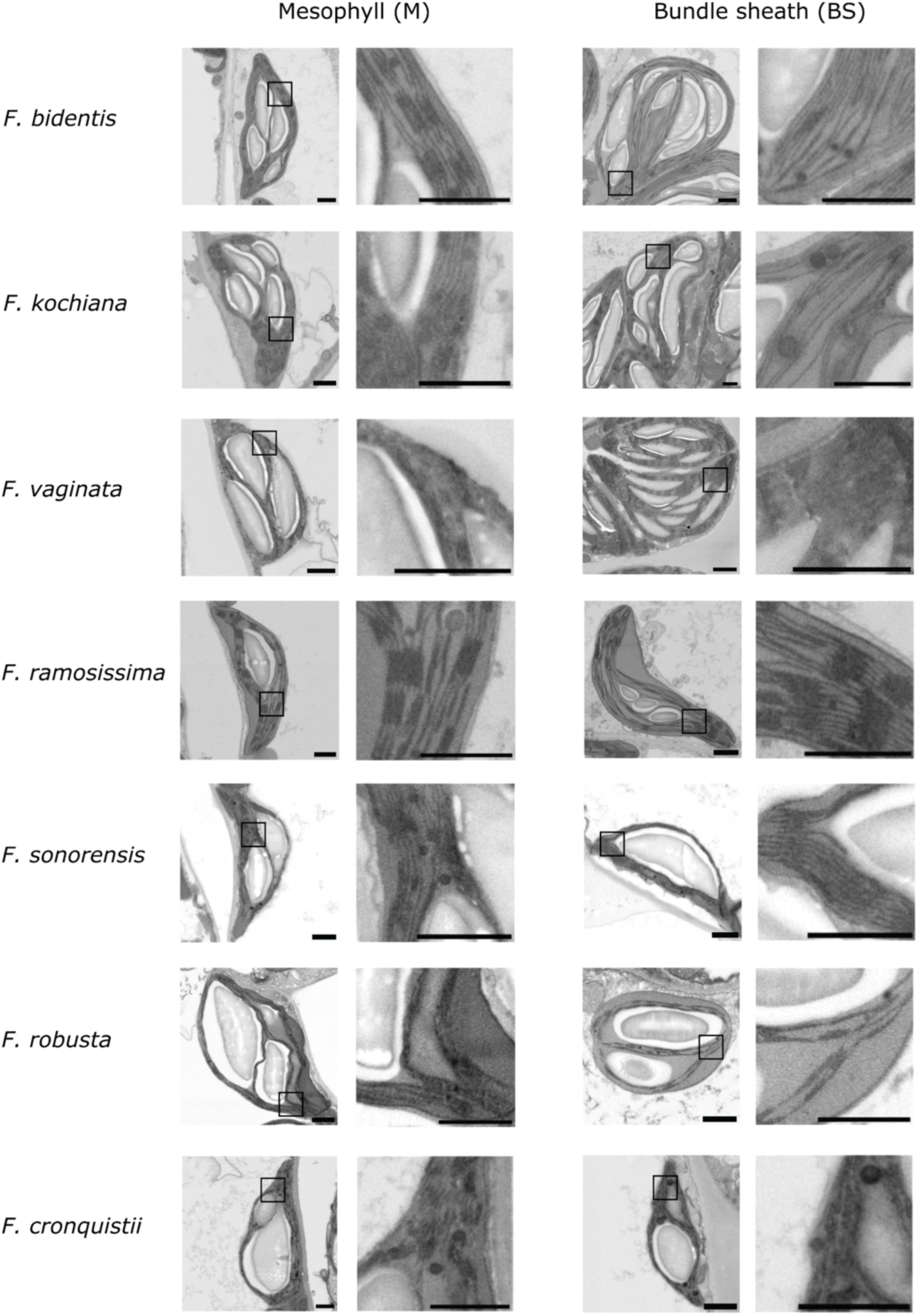
Representative excerpts from 2D high-resolution (12,000x) scanning electron microscopy maps showing chloroplast ultrastructure in mesophyll (M) and bundle sheath (BS) cells across the *Flaveria* species. For each chloroplast image, a corresponding zoomed-in view is shown. The zoomed-in area is indicated by a black box. Bars = 1 μm

**Supplemental Figure S6:**
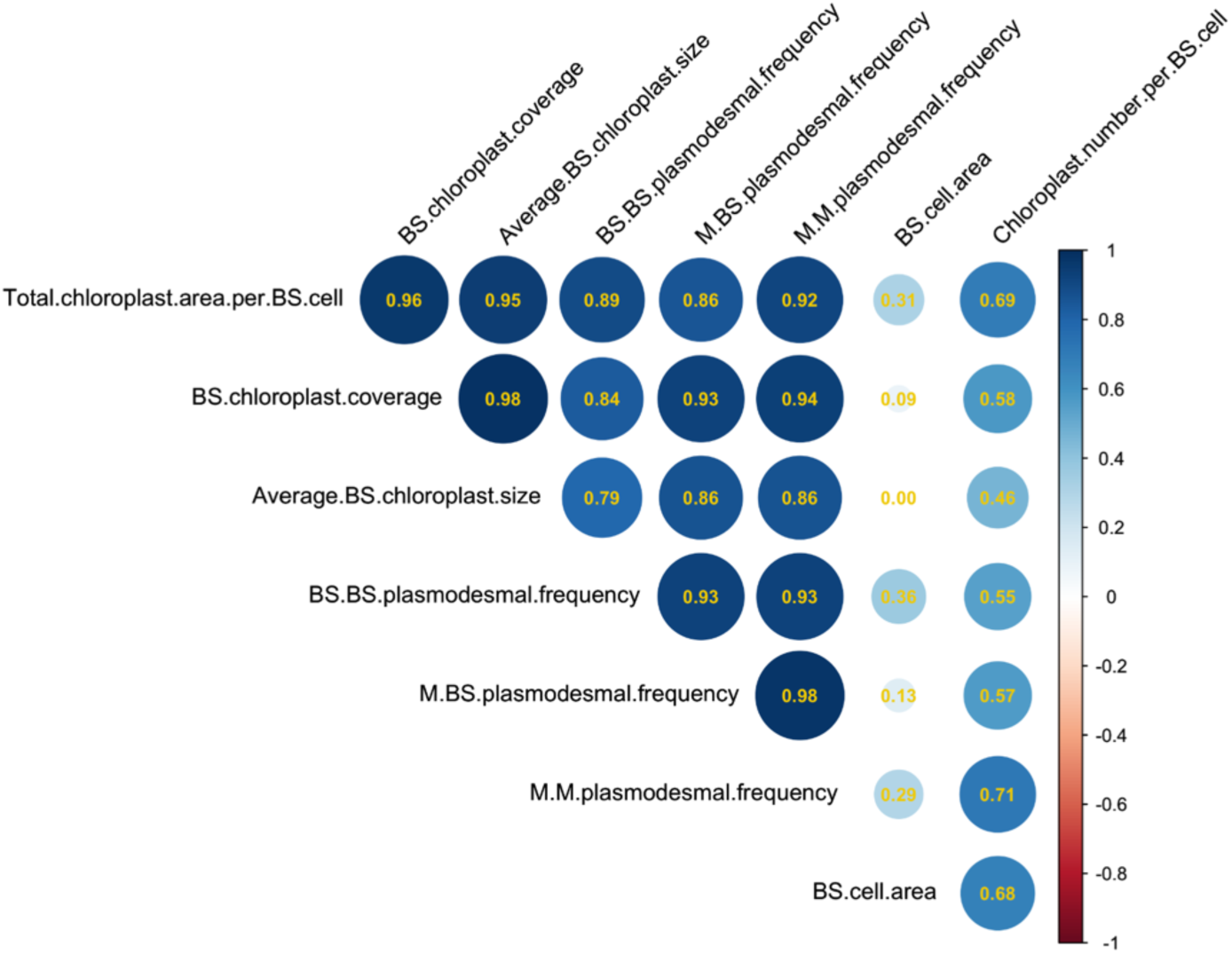
Correlation analysis for plasmodesmal frequencies (at mesophyll-bundle sheath (M-BS), mesophyll-mesophyll (M-M), and bundle sheath-bundle sheath (BS-BS) cell interfaces), and bundle sheath (BS) cell and chloroplast-related anatomical traits along the C_3_-to-C_4_ evolutionary gradient in *Flaveria*. The correlation coefficients were derived by Pearson correlation analysis and presented in each circle in yellow. The matrix is hierarchically clustered and reordered based on Pearson correlation coefficients. Correlation strength is also colour-coded, with blue representing positive correlation, white indicating no correlation, and red indicating negative correlation.

## Notes

### Competing Interest Statement

The authors have declared no competing interest.

## References

Alenazi AS, Pereira L, Christin P, Osborne CP, Dunning LT. 2024. Identifying genomic regions associated with C4 photosynthetic activity and leaf anatomy in *Alloteropsis semialata*. New Phytologist 243: 1698–1710.

Azim MF, Burch-Smith TM. 2020. Organelles-nucleus-plasmodesmata signaling (ONPS): an update on its roles in plant physiology, metabolism and stress responses. Current Opinion in Plant Biology 58: 48–59.

Bayer EM, Benitez-Alfonso Y. 2024. Plasmodesmata: Channels Under Pressure. Annual Review of Plant Biology 75: 291–317.

Benitez-Alfonso Y, Cilia M, Roman AS, Thomas C, Maule A, Hearn S, Jackson D. 2009. Control of *Arabidopsis* meristem development by thioredoxin-dependent regulation of intercellular transport. Proceedings of the National Academy of Sciences 106: 3615–3620.

Borghi GL, Arrivault S, Günther M, Barbosa Medeiros D, Dell’Aversana E, Fusco GM, Carillo P, Ludwig M, Fernie AR, Lunn JE, et al. 2022. Metabolic profiles in C3, C3–C4 intermediate, C4-like, and C4 species in the genus *Flaveria* (A Braeutigam, Ed.). Journal of Experimental Botany 73: 1581–1601.

Botha CEJ. 1992. Plasmodesmatal distribution, structure and frequency in relation to assimilation in C3 and C4 grasses in southern Africa. Planta 187.

Botha CEJ, Cross RHM, Van Bel AJE, Peter CI. 2000. Phloem loading in the sucrose-export-defective (SXD-1) mutant maize is limited by callose deposition at plasmodesmata in bundle sheath—vascular parenchyma interface. Protoplasma 214: 65–72.

Botha CEJ, Hartley BJ, Cross RHM. 1993. The Ultrastructure and Computer-enhanced Digital Image Analysis of Plasmodesmata at the Kranz Mesophyll-Bundle Sheath Interface of Themeda triandra var. imberbis (Retz) A. Camus in Conventionally-fixed Blades. Annals of Botany 72: 255–261.

Brown NJ, Parsley K, Hibberd JM. 2005. The future of C4 research – maize, Flaveria or Cleome? Trends in Plant Science 10: 215–221.

Burch-Smith TM, Zambryski PC. 2010. Loss of INCREASED SIZE EXCLUSION LIMIT (ISE)1 or ISE2 Increases the Formation of Secondary Plasmodesmata. Current Biology 20: 989–993.

Burch-Smith TM, Zambryski PC. 2012. Plasmodesmata Paradigm Shift: Regulation from Without Versus Within. Annual Review of Plant Biology 63: 239–260.

Danila FR, Quick WP, White RG, Furbank RT, Von Caemmerer S. 2016. The Metabolite Pathway between Bundle Sheath and Mesophyll: Quantification of Plasmodesmata in Leaves of C 3 and C 4 Monocots. The Plant Cell 28: 1461–1471.

Danila FR, Quick WP, White RG, Kelly S, Von Caemmerer S, Furbank RT. 2018. Multiple mechanisms for enhanced plasmodesmata density in disparate subtypes of C4 grasses. Journal of Experimental Botany 69: 1135–1145.

Danila FR, Quick WP, White RG, Von Caemmerer S, Furbank RT. 2019. Response of plasmodesmata formation in leaves of C4 grasses to growth irradiance. Plant, Cell & Environment 42: 2482–2494.

Drincovich MF, Casati P, Andreo CS, Chessin SJ, Franceschi VR, Edwards GE, Ku MSB. 1998. Evolution of C4 Photosynthesis in *Flaveria* Species1. Plant Physiology 117: 733–744.

Edwards GE, Ku MSB. 1987. Biochemistry of C3-C4 intermediates. In: Hatch M, Boardman N, eds. The Biochemistry of Plants. London: Academic Press, 275–325.

Evert RF, Eschrich W, Heyser W. 1977. Distribution and structure of the plasmodesmata in mesophyll and bundle-sheath cells of Zea mays L. Planta 136: 77–89.

Faulkner C, Akman OE, Bell K, Jeffree C, Oparka K. 2008. Peeking into Pit Fields: A Multiple Twinning Model of Secondary Plasmodesmata Formation in Tobacco. The Plant Cell 20: 1504–1518.

Ganusova EE, Reagan BC, Fernandez JC, Azim MF, Sankoh AF, Freeman KM, McCray TN, Patterson K, Kim C, Burch-Smith TM. 2020. Chloroplast-to-nucleus retrograde signalling controls intercellular trafficking via plasmodesmata formation. Philosophical Transactions of the Royal Society B: Biological Sciences 375: 20190408.

Gowik U, Bräutigam A, Weber KL, Weber APM, Westhoff P. 2011. Evolution of C4 Photosynthesis in the Genus *Flaveria* : How Many and Which Genes Does It Take to Make C4? The Plant Cell 23: 2087–2105.

Hatch MD. 1987. C4 photosynthesis: a unique mena of modified biochemistry, anatomy and ultrastructure. Biochimica et Biophysica Acta 895: 81–106.

Keerberg O, Pärnik T, Ivanova H, Bassüner B, Bauwe H. 2014. C2 photosynthesis generates about 3-fold elevated leaf CO2 levels in the C3–C4 intermediate species Flaveria pubescens. Journal of Experimental Botany 65: 3649–3656.

Khoshravesh R, Stata M, Busch FA, Saladié M, Castelli JM, Dakin N, Hattersley PW, Macfarlane TD, Sage RF, Ludwig M, et al. 2020. The Evolutionary Origin of C4 Photosynthesis in the Grass Subtribe Neurachninae. Plant Physiology 182: 566–583.

Khoshravesh R, Stinson CR, Stata M, Busch FA, Sage RF, Ludwig M, Sage TL. 2016. C3 –C4 intermediacy in grasses: organelle enrichment and distribution, glycine decarboxylase expression, and the rise of C2 photosynthesis. Journal of Experimental Botany 67: 3065–3078.

Kobayashi K, Otegui MS, Krishnakumar S, Mindrinos M, Zambryski P. 2007. *INCREASED SIZE EXCLUSION LIMIT2* Encodes a Putative DEVH Box RNA Helicase Involved in Plasmodesmata Function during *Arabidopsis* Embryogenesis. The Plant Cell 19: 1885–1897.

Ku MSB, Monson RK, Littlejohn RO, Nakamoto H, Fisher DB, Edwards G. 1983. Photosynthetic Characteristics of C₃-C₄ Intermediate Flaveria Species: I. Leaf Anatomy, Photosynthetic Responses to O₂ and CO₂ and Activities of Key Enzymes in the C₃ and C₄ Pathways. Plant Physiology 71: 944–948.

Kümpers BMC, Burgess SJ, Reyna-Llorens I, Smith-Unna R, Boursnell C, Hibberd JM. 2017. Shared characteristics underpinning C4 leaf maturation derived from analysis of multiple C3 and C4 species of *Flaveria*. Journal of Experimental Botany 68: 177–189.

Lauterbach M, Bräutigam A, Clayton H, Saladié M, Rolland V, Macfarlane TD, Weber APM, Ludwig M. 2024. Leaf transcriptomes from C3, C3-C4 intermediate, and C4 *Neurachne* species give insights into C4 photosynthesis evolution. Plant Physiology 197: kiae424.

Li ZP, Paterlini A, Glavier M, Bayer EM. 2021. Intercellular trafficking via plasmodesmata: molecular layers of complexity. Cellular and Molecular Life Sciences 78: 799–816.

Lundgren MR. 2020. C2 photosynthesis: a promising route towards crop improvement? New Phytologist 228: 1734–1740.

Lundgren MR, Christin P, Escobar EG, Ripley BS, Besnard G, Long CM, Hattersley PW, Ellis RP, Leegood RC, Osborne CP. 2016. Evolutionary implications of C3 –C4 intermediates in the grass *Alloteropsis semialata*. Plant, Cell & Environment 39: 1874–1885.

Lyu M-JA, Gowik U, Kelly S, Covshoff S, Mallmann J, Westhoff P, Hibberd JM, Stata M, Sage RF, Lu H, et al. 2015. RNA-Seq based phylogeny recapitulates previous phylogeny of the genus Flaveria (Asteraceae) with some modifications. BMC Evolutionary Biology 15: 116.

McKown AD, Dengler NG. 2007. Key innovations in the evolution of Kranz anatomy and C4 vein pattern in *Flaveria* (Asteraceae). American Journal of Botany 94: 382–399.

Mercado MA, Studer AJ. 2022. Meeting in the Middle: Lessons and Opportunities from Studying C3 -C4 Intermediates. Annual Review of Plant Biology 73: 43–65.

Monson RK, Moore BD. 1989. On the significance of C3 —C4 intermediate photosynthesis to the evolution of C4 photosynthesis. Plant, Cell & Environment 12: 689–699.

Muhaidat R, Sage TL, Frohlich MW, Dengler NG, Sage RF. 2011. Characterization of C3 –C4 intermediate species in the genus *Heliotropium* L. (Boraginaceae): anatomy, ultrastructure and enzyme activity. *Plant*, Cell & Environment 34: 1723–1736.

Osmond C. 1971. Metabolite Transport in C4 Photosynthesis. Australian Journal of Biological Sciences 24: 159.

Otero S, Helariutta Y, Benitez-Alfonso Y. 2016. Symplastic communication in organ formation and tissue patterning. Current Opinion in Plant Biology 29: 21–28.

Provencher LM, Miao L, Sinha N, Lucas WJ. 2001. Sucrose Export Defective1 Encodes a Novel Protein Implicated in Chloroplast-to-Nucleus Signaling. The Plant Cell 13: 1127–1141.

Roberts AG, Oparka KJ. 2003. Plasmodesmata and the control of symplastic transport. Plant, Cell & Environment 26: 103–124.

Sage RF. 2004. The evolution of C4 photosynthesis. New Phytologist 161: 341–370.

Sage TL, Busch FA, Johnson DC, Friesen PC, Stinson CR, Stata M, Sultmanis S, Rahman BA, Rawsthorne S, Sage RF. 2013. Initial Events during the Evolution of C4 Photosynthesis in C3 Species of *Flaveria*. Plant Physiology 163: 1266–1276.

Sage RF, Christin P-A, Edwards EJ. 2011. The C4 plant lineages of planet Earth. Journal of Experimental Botany 62: 3155–3169.

Schlüter U, Bouvier JW, Guerreiro R, Malisic M, Kontny C, Westhoff P, Stich B, Weber APM. 2023. *Brassicaceae* display variation in efficiency of photorespiratory carbon-recapturing mechanisms (J Lunn, Ed.). Journal of Experimental Botany 74: 6631–6649.

Schlüter U, Bräutigam A, Gowik U, Melzer M, Christin P-A, Kurz S, Mettler-Altmann T, Weber AP. 2017. Photosynthesis in C3 –C4 intermediate *Moricandia* species. Journal of Experimental Botany 68: 191–206.

Schreier TB, Müller KH, Eicke S, Faulkner C, Zeeman SC, Hibberd JM. 2024. Plasmodesmal connectivity in C 4 *Gynandropsis gynandra* is induced by light and dependent on photosynthesis. New Phytologist 241: 298–313.

Schulze S, Mallmann J, Burscheidt J, Koczor M, Streubel M, Bauwe H, Gowik U, Westhoff P. 2013. Evolution of C4 Photosynthesis in the Genus Flaveria: Establishment of a Photorespiratory CO2 Pump. The Plant Cell 25: 2522–2535.

Schüssler C, Freitag H, Koteyeva N, Schmidt D, Edwards G, Voznesenskaya E, Kadereit G. 2017. Molecular phylogeny and forms of photosynthesis in tribe Salsoleae (Chenopodiaceae). Journal of Experimental Botany 68: 207–223.

Sedelnikova OV, Hughes TE, Langdale JA. 2018. Understanding the Genetic Basis of C 4 Kranz Anatomy with a View to Engineering C 3 Crops. Annual Review of Genetics 52: 249–270.

Stata M, Sage TL, Hoffmann N, Covshoff S, Ka-Shu Wong G, Sage RF. 2016. Mesophyll Chloroplast Investment in C3 , C4 and C2 Species of the Genus *Flaveria*. Plant and Cell Physiology 57: 904–918.

Stata M, Sage TL, Rennie TD, Khoshravesh R, Sultmanis S, Khaikin Y, Ludwig M, Sage RF. 2014. Mesophyll cells of C4 plants have fewer chloroplasts than those of closely related C3 plants. *Plant*, Cell & Environment 37: 2587–2600.

Stitt M, Heldt HW. 1985. Control of photosynthetic sucrose synthesis by fructose-2,6-bisphosphate. Planta 164: 179–188.

Stonebloom S, Burch-Smith T, Kim I, Meinke D, Mindrinos M, Zambryski P. 2009. Loss of the plant DEAD-box protein ISE1 leads to defective mitochondria and increased cell-to-cell transport via plasmodesmata. Proceedings of the National Academy of Sciences 106: 17229–17234.

Tee EE, Faulkner C. 2024. Plasmodesmata and intercellular molecular traffic control. New Phytologist 243: 32–47.

Walker BJ, VanLoocke A, Bernacchi CJ, Ort DR. 2016. The Costs of Photorespiration to Food Production Now and in the Future. Annual Review of Plant Biology 67: 107–129.

Walsh CA, Bräutigam A, Roberts MR, Lundgren MR. 2023. Evolutionary implications of C2 photosynthesis: how complex biochemical trade-offs may limit C4 evolution (D Ort, Ed.). Journal of Experimental Botany 74: 707–722.

Wang P, Khoshravesh R, Karki S, Tapia R, Balahadia CP, Bandyopadhyay A, Quick WP, Furbank R, Sage TL, Langdale JA. 2017. Re-creation of a Key Step in the Evolutionary Switch from C3 to C4 Leaf Anatomy. Current Biology 27: 3278–3287.e6.

Zhao Y-Y, Lyu MA, Miao F, Chen G, Zhu X-G. 2022. The evolution of stomatal traits along the trajectory toward C4 photosynthesis. Plant Physiology 190: 441–458.

